# Mechanisms of octopus arm search behavior without visual feedback

**DOI:** 10.1101/2023.03.13.532148

**Authors:** Dominic M. Sivitilli, Terrell Strong, Willem Weertman, Joseph Ullmann, Joshua R. Smith, David H. Gire

## Abstract

The octopus coordinates multiple, highly flexible arms with the support of a complex distributed nervous system. The octopus’s suckers, staggered along each arm, are employed in a wide range of behaviors. Many of these behaviors, such as foraging in visually occluded spaces, are executed under conditions of limited or absent visual feedback. In coordinating unseen limbs with seemingly infinite degrees of freedom across a variety of adaptive behaviors, the octopus appears to have solved a significant control problem facing the field of soft-bodied robotics. To study the strategies that the octopus uses to find and capture prey within unseen spaces, we designed and 3D printed visually occluded foraging tasks and tracked arm motion as the octopus attempted to find and retrieve a food reward. By varying the location of the food reward within these tasks, we can characterize how the arms and suckers adapt to their environment to find and capture prey. We compared these results to simulated experimental conditions performed by a model octopus arm to isolate the primary mechanisms driving our experimental observations. We found that the octopus relies on a contact-based search strategy that emerges from local sucker coordination to simplify the control of its soft, highly flexible limbs.

## Introduction

The octopus employs its eight highly flexible arms across a range of behaviors, including foraging, exploration, manipulation, and locomotion. Each arm has hundreds of suckers staggered along its length. These suckers bear a complex chemotactile system (Graziadei, 1964; Graziadei & Gagne, 1976; Graziadei, 1965a; van Giesen et al., 2020) and are the primary appendages used by the octopus to interact with its environment (Packard et al., 1988). Most of the octopus’s nervous system is distributed into its arms and suckers (Young, 1971), and locally controls behavior with minimal feedback from the brain. Much of the arm’s behavioral repertoire therefore remains intact when the arm is denervated from the brain (Gutfreund et al., 2006; Altman, 1968; Rowell, 1963; Zullo, 2011; Sumbre et al., 2001).

The brain and arms communicate over a limited neural bandwidth (Young, 1965) and a great deal of information within the arms, such as proprioceptive information from stretch receptors in the arm musculature, has been found to not be communicated to the brain (Rowell, 1966; Wells & Wells, 1957; Wells, 1964; Graziadei, 1965b). With a limited representation of the configuration of the arms and suckers, the brain is likewise limited in its ability to generate a detailed motor plan. Instead, it has been suggested that the brain sends out a generalized motor plan to the arms, where it activates peripheral neural circuitry to locally control behavior under the direction of the suckers’ complex chemotactile system (Zullo et al., 2009; Zullo et al., 2019). This reliance on local control in place of centralized planning means that the brain is also limited in its ability to recall the details of arm behavior (Wells & Wells, 1957; Wells, 1964).

Possibly due to the lack of sensory feedback from the arms, reaching behavior relies on a feedforward activation mechanism (Gutfreund et al., 1998; Sumbre et al., 2001), which gives the arm a ballistic kinematic profile as it extends toward its target. This suggests that the brain only takes into account the vertical and horizontal angle (yaw and pitch) of a target and activates reaching behavior in this direction without any further modification of the behavior following its activation. Where this information is sufficient for the retrieval of a reward, the octopus shows an improved performance over time without visual feedback (Gutnick et al., 2020). Beyond these limited parameters that the brain is able to control and recall, the suckers and their local nervous system likely provides the necessary sensory-motor feedback to adapt behavior to the environment.

In tasks where visual feedback is absent and feedforward activation is inadequate for the retrieval of a reward, the octopus must rely primarily on the chemotactile system of the suckers in the generation and modification of behavior. Characterizing the strategies that are employed by the arms and suckers to search for and retrieve a reward in the absence of visual feedback can therefore help resolve how the chemotactile system is used to control the arm’s extreme flexibility. Here, we investigate the strategies used by the Pacific red octopus (*Octopus rubescens*) when searching for and retrieving a food reward from a visually occluded foraging task space.

Previous studies have described a locally controlled behavior within the arm which we refer to here as sucker recruitment. During this behavior, which can be elicited when the arm is denervated from the brain, sensory input to a sucker results in the neighboring suckers bending toward the source of this sensory input (Gutfreund et al., 2006; Altman, 1968; Rowell, 1963; Zullo, 2011). This same effect can then be elicited in these neighbors and cause a propagating wave of recruitment down the arm. This recruitment mechanism can serve as an effective strategy for object handling and prey capture, and could likewise serve as a potential foraging strategy by allowing the arm to adapt to the shape of complex surfaces in search of prey hidden within unseen crevices. This form of surface conformation could further provide a means for the octopus to simplify control of its movement by providing constraints on the arm’s vast degrees of freedom. The reliance on contact would be consistent with a force control (rather than position or velocity control) strategy by the octopus.

We therefore predict that if the octopus reaches its arm through a narrow entrance into an open space, sucker contact with the entrance will initiate a recruitment signal which will attract the arm toward the surfaces of the space shared by the entrance. We also predict that the arm will preferentially search along concave edges and vertices where multiple surfaces meet, as these contours could serve to confine the arm’s range of possible configurations and thereby simplify behavior.

To investigate the role of sucker recruitment in generating our experimental observations, we created a simple computational model of the octopus arm and subjected it to simulated experimental conditions that precisely matched our 3D-printed experimental task space. This model included a recruitment mechanism that could be activated between serially joined segments, which operated in parallel to random motion generated within the joints. This random motion was amplified at the proximal-most joint of the model, which corresponded to the section of the octopus’s arm reaching through the task entrance.

## Methods

Subjects for this study included eight Pacific red octopuses (*Octopus rubescens*) collected using SCUBA from the Puget Sound. All animals were collected under an approved permit through the Washington Department of Fish and Wildlife. Animals were kept in tanks that ranged in volume from 3 to 40 gallons (roughly 10-150 liters) depending on the size of the animal, tank availability, and enrichment schedule. During the course of the study, the subjects were offered food (scallop or shrimp meat) equal to 1-2% of their mass daily as the reward for the task. All animals were enriched and handled regularly during the course of the experiment. All experiments were carried out in accordance with a protocol approved by the University of Washington Institutional Animal Care and Use Committee (IACUC).

Figure 1 outlines our experimental design. The foraging task and its interior task space were created using computer-aided design (CAD) software and 3D-printed with white polylactic acid (PLA) filament. This task was secured to the side of a glass or transparent acrylic tank using magnets, such that the task space was visible to a camera outside the tank, but visually occluded to the octopus. At the top of the task was a circular entrance large enough for a single arm to fit through. This entrance was at the bottom of a larger opening designed to guide the octopus’s arms toward the entrance. The task space was shaped as a half-cylinder that connected the entrance on its upward-facing round side to a rectangular box on its downward-facing flat side. The box and half cylinder had the same depth along the axis between the tank and the camera. The task was printed in two sizes depending on the weight of the subject. Octopuses over 80g were assigned the large task and octopuses under 60g were assigned the small task, which was 80% the size of the large task. Octopuses between 60-80g were randomly assigned to either version. A light was used to illuminate the task from behind to maximize the contrast between the task space features and the octopus’s arm. An acrylic barrier prevented the octopus from creating shadow artifacts by reaching between the task and the light. Prior to the start of trials, the octopuses were trained to approach the entrance of the task for a food reward while the interior was blocked off.

**Figure 1.**
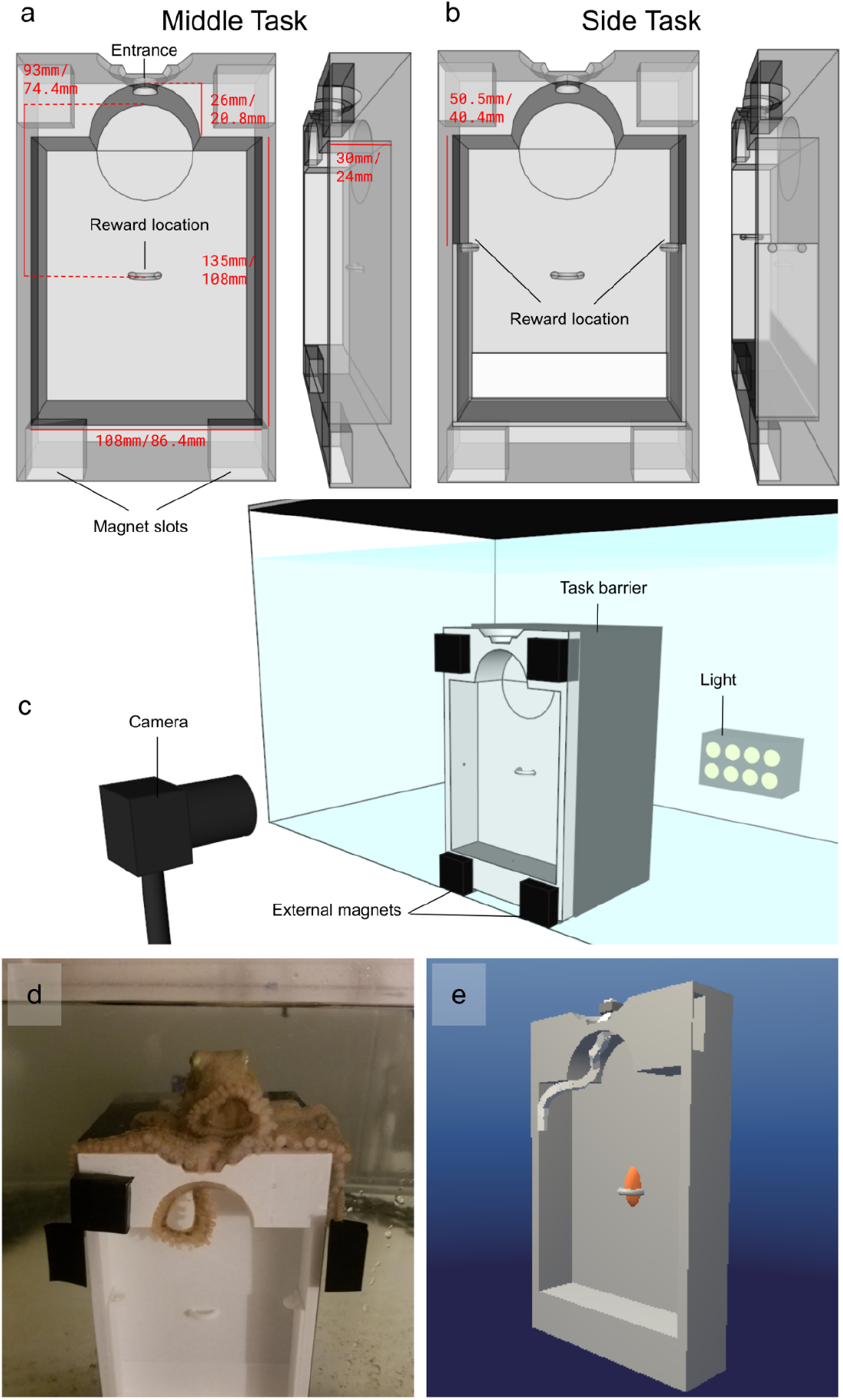
Experimental Design. (a) Middle and (b) Side task CAD design with dimensions of both large and small task versions displayed. (c) Experimental setup. (d) Octopus during attempt. (e) Model arm during simulated task.

For one condition (“Middle” ; n = 4 males), the food reward was secured in the center of the box’s largest face, and in the other condition (“Side”), the reward was secured to either the left (n = 2; one male and one female) or right wall (n = 2; one male and one female) of the box close to the concave edge between the wall and the box’s largest face. Video recording failed for the first trial of one subject in the side condition, so this trial was not included in the analysis. The reward was equally distant from the entrance in both conditions.

The octopuses were given as many opportunities as needed to retrieve the reward six times from the task. Each of these six trials included the total time the octopus spent searching the task space until the reward was found. The time between when an arm entered the task space and when it either found the reward or left unsuccessfully was considered an attempt. Trials often consisted of multiple failed attempts before a successful attempt was made. An example of a successful attempt for the middle and side task can be seen in Supplemental Video 2 and Supplemental Video 3, respectively. Arm behavior was recorded at 250 FPS with a CMOS camera (Imaging Source, model DMK 37BUX287) and custom-written software in LabVIEW (National Instruments). Arms were tracked using DeepLabCut pose estimation software (Mathis et al., 2018) (see Supplemental Figure 1), and analysis was performed using custom routines written in Python.

The computational model was developed using the Unity game engine as a chain of rigid body segments connected by joints with three degrees of freedom (yaw, pitch, and roll). The mechanics of the arm were written as routines in C# and performed two primary functions. One of these was to simulate sucker recruitment in response to contact with environmental features. Upon touching a surface, torque was applied to the neighboring segments to rotate them in the direction of the feature. This caused the arm to conform to surface features as each segment propagated this recruitment signal in response to touching the feature. The second function was to generate random motion of the arm. This took the form of a random intensity of torque from a range applied to each segment in a random direction along the yaw and pitch of its proximal joint after a randomly selected delay between two and three seconds. The intensity for the proximal-most joint, connecting the segment reaching through the entrance to a stationary segment just outside the entrance, was 20x that of the other joints. The segments were also subjected to a drag force to simulate motion through liquid. The task was simulated by importing the task CAD file into Unity. The arm began each trial in a straight vertical pose extending down from the entrance. A simulated trial was completed when the arm made contact with the reward, which approximated the size and shape of the food rewards from the experiments. An example of a simulated trial for the middle task and both versions of the side task are shown in Supplemental Video 4.

## Results

Performance for the observed data was assessed based on the time taken to reach the reward over the six trials and between the two conditions (time to success, Figure 2). There was no improvement in performance across the six trials for either condition, but there was notably consistently better performance on the side task across trials. When trial averages were compared between conditions, the octopuses performed significantly better on the side task (Mann-Whitney *U*, p < 0.05).

**Figure 2.**
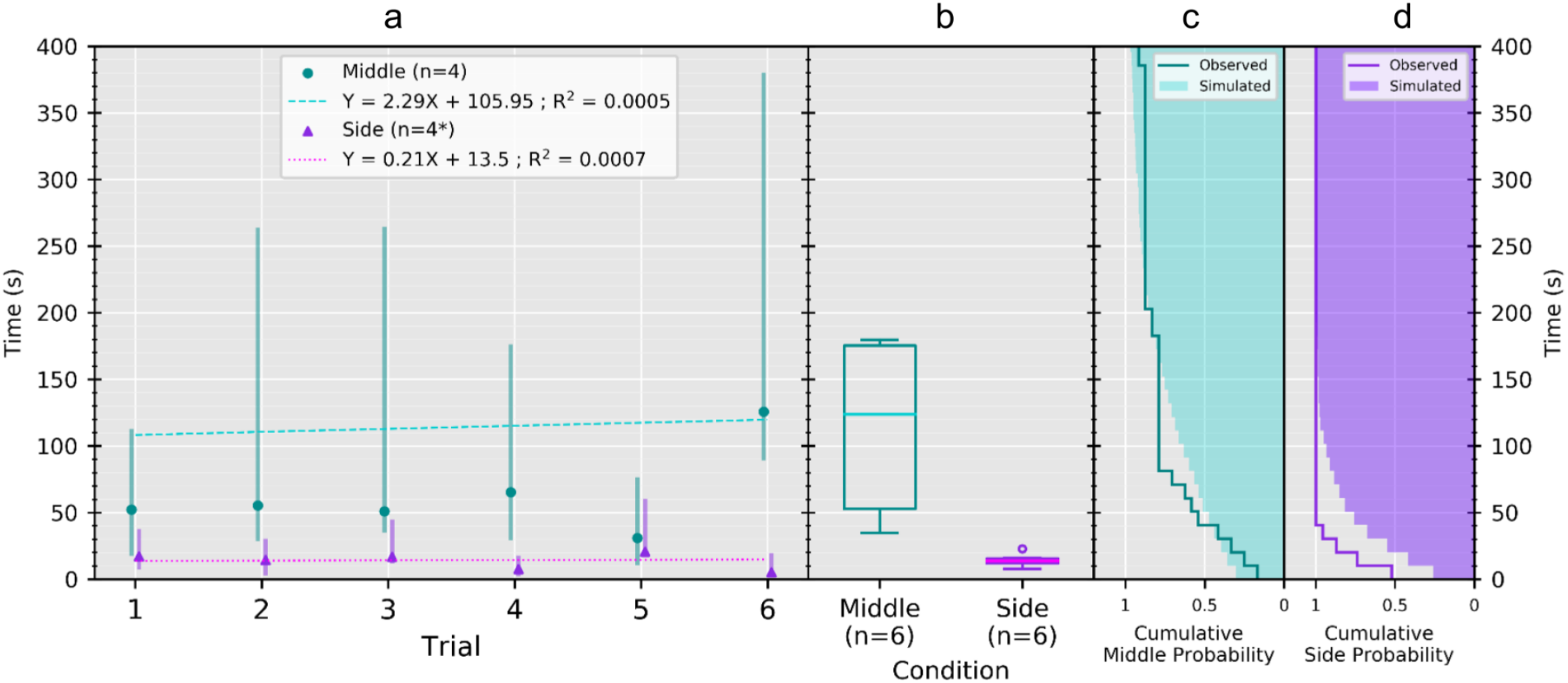
Performance as time to success. (a) Performance over trials (as median and interquartile range), and regression showing no improvement. (b) Performance between conditions as trial averages, showing significantly better performance for the side task (Mann-Whitney *U*; p < 0.05). (c) Middle cumulative probability distributions of observed and simulated performance, showing shared distribution (K-S; p = 0.21). The maximum observed and simulated trial duration lasted 604.34 and 916.56 seconds, respectively. (d) Side cumulative probability distributions of observed and simulated performance, showing discrete distributions (K-S; p = 0.01). *Video recording failed during the first trial of one subject in the side condition, so this trial was not included in the analysis.

Three kinematic measures were used to characterize arm behavior during the octopus’s attempts to find the food reward: arm segment speed, curvature, and wall proximity (Figure 3). All three of these measures showed a recurring behavioral profile represented by a distally oriented wave of deceleration, curvature and movement toward the wall, suggesting that the proximal arm leads the initiation of this behavior. The wave appeared to be a result of a distally propagating wave of sucker recruitment initiated and continuously regenerated by sucker contact with the task space features. This resulted in the arm conforming to the shape of these features, which was likewise reflected in arm occupancy across animals and trials (Figure 4), showing the arms spending most of their time conforming to the sides of the task during the experiment.

**Figure 3.**
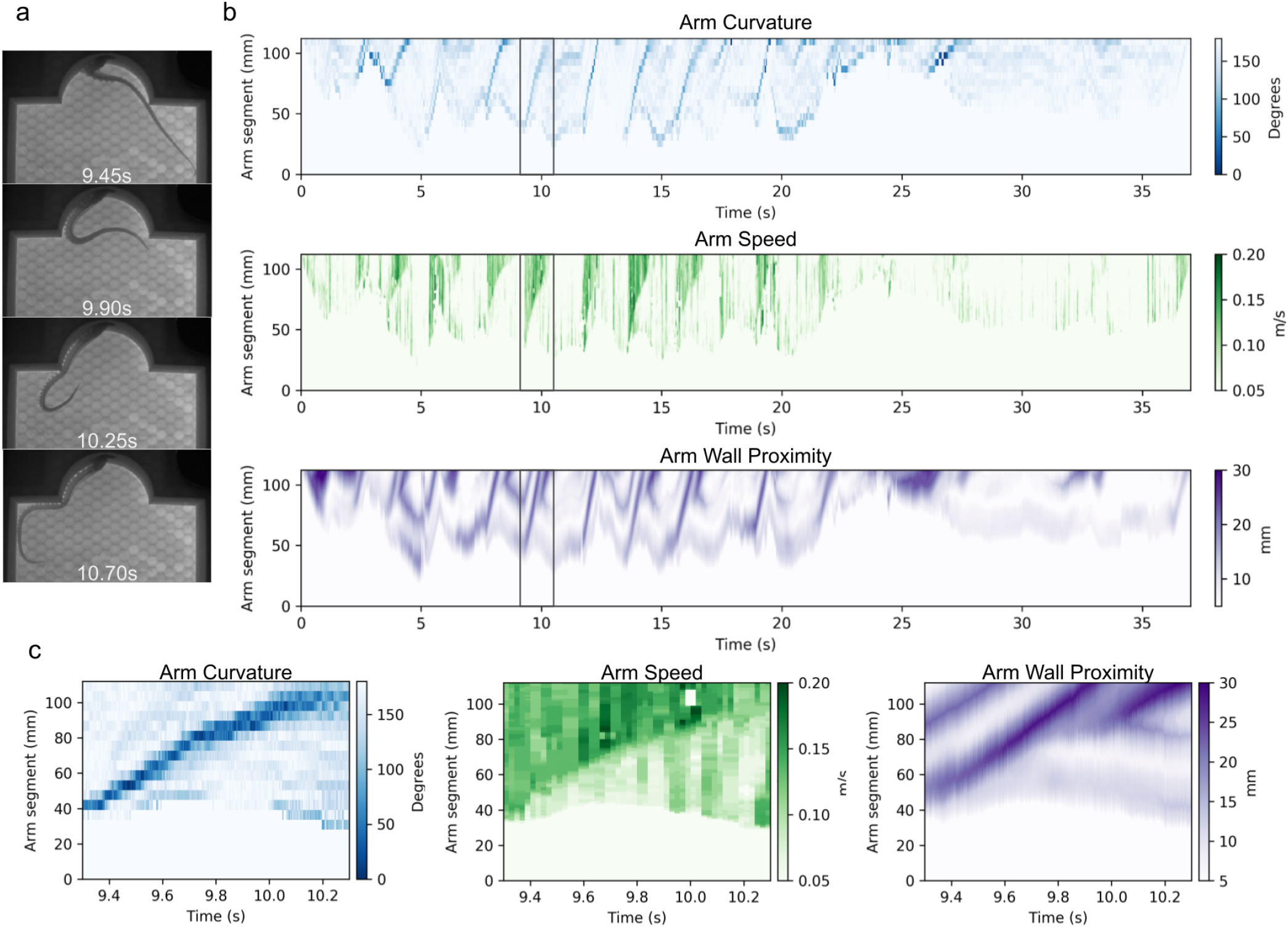
(b) Example timeplots representing arm curvature (top), speed (middle), and wall proximity (bottom) during a middle attempt (shown in Supplemental Video 1) using the large version of the task. Horizontal axes represent time and vertical axes represent segments of the arm with the arm tip at the top. (a & c) Recurring kinematic profile of a propagating wave of curvature, deceleration, and movement toward the wall. Note that curvature is represented here by the angle between segments in degrees, such that the lower the value the greater the curvature.

**Figure 4.**
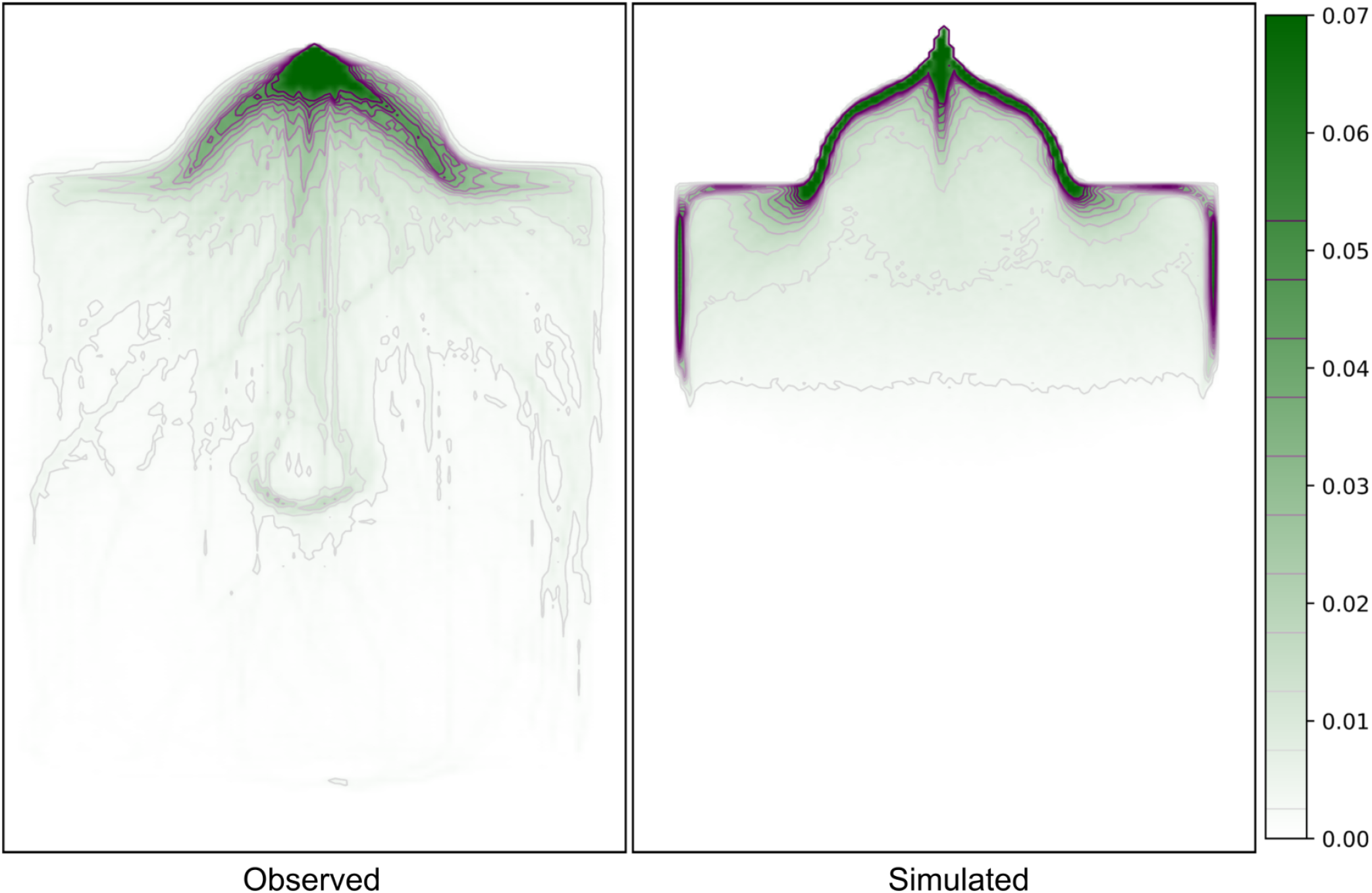
Most commonly occupied areas of the task for (left, “Observed”) the observed arms as proportion of total time across all trials and all animals (equally weighted) and for (right, “Simulated”) the model over 1,000 seconds of a simulation without a reward to find. Both the arms of the observed data and model in the simulated data were represented by a single pixel-width line to calculate these plots.

Using the model, we ran 1,000 simulations for each condition. We then compared the performance distribution for each simulated condition to our observed performance data using a Kolmogorov-Smirnov (K-S) test. Results suggest that the observed and simulated arms share a distribution for the middle task (p = 0.21), but not for the side task (p = 0.01). This shows that the model accurately simulated performance when the reward was found in the center of the task space. While the observed and simulated distribution for the side task appear to share a profile, the experimentally observed arms in this case acquire the reward consistently faster.

We ran a single simulation for 1,000 seconds to characterize the preferred occupancy of the model (see Figure 4). No reward was included in this simulation to prevent the model from being reset to its starting pose upon finding the reward. Like the experimentally-observed arms, the model preferred to conform to the shape of the task space features, which suggests that the model accurately simulated arm occupancy.

## Discussion

To describe the control problem that the octopus faces, we will introduce the concept ‘configuration space’ used in the field of robotics to represent the multidimensional space of all possible configurations of a robot, where each axis represents a degree of freedom of the robot’s joint angles. The more joints a robot has, the higher the dimensionality of its configuration space. With the octopus arm able to bend at any continuous point along its length in any direction, the octopus’s space of possible configurations is immeasurably large (Kennedy et al., 2020). This is all the more striking given that the octopus employs its arms across a wide range of behaviors. By using limbs with effectively infinite degrees of freedom across a variety of adaptive behaviors, the octopus has solved a significant control problem facing the field of soft-bodied robotics. If we can identify and characterize the biomechanical properties and control strategies the octopus employs to do this, we can implement these elements in the development of soft-robotic limbs with a similar range of capabilities.

Here we investigated the strategies that the octopus uses to search for prey without visual feedback, which is often the case when foraging within crevices, under rocks, or in conditions of low visibility. In these cases the role of vision is removed, and without a central representation of arm configuration, the octopus must rely primarily on the chemotactile feedback received by the suckers and the behaviors locally controlled within the arm’s nerve cord.

To identify the mechanisms underlying these behaviors, our approach is to program these mechanisms into our computational model, subject the model to simulated experimental conditions, then compare the model’s behavior to that of the octopus arm. The more accurately the mechanism is replicated within the model and the larger role the mechanism plays in the arm’s behavior, the more closely the model would reasonably match the experimental data.

The prevalence of sucker recruitment in behavior with and without a connection to the brain made it a compelling mechanism to investigate within this modeling framework. As this mechanism alone would cause the model to stick motionless against a surface, we paired it with random motion generated within the segments. This motion was amplified in the proximal-most joint to simulate the proximally-led, distally oriented recruitment signal observed from the kinematic data.

Together, these mechanisms resulted in a pattern of the proximal segment “casting” the arm in a random direction, where the rest of the segments then serially conformed to the task space features. Between these larger casting motions, the random motion of the distal segments caused the arm to move laterally across the surface.

The performance and occupancy of our experimental results suggest that the arms preferentially conformed to the shape of the task space’s surface features, particularly along its concave edges, and that this behavior led to greater performance when the reward was found on the side.

To isolate sucker recruitment as the underlying mechanism of this behavior, we compared the performance distributions of our experimental data with those of our simulations. This comparison indicated that the observed and simulated arm share a distribution for the middle task, but not for the side task. While observed and simulated distributions shared a similar profile for the side task, the observed arms found the reward significantly faster. There are a number of possible reasons that could explain why the octopus performed better than the model that highlight the model’s limitations.

While the random motion generated a performance and occupancy profile that resembled the observed arms, the octopus likely employs a more systematic search strategy. For example, when conforming to a surface, the octopus arm’s lateral movement is most likely not equally probable in both directions. Also, rather than generating larger proximal movements within a random interval of time, this kind of behavior probably occurs after the arm exhausts the novelty of a searched area. It is reasonable to suggest that these two factors are not as limiting when the reward was found in the middle, which may have accounted for the closer resemblance in performance between the model and the observed arm for the middle task.

A systematic search strategy of the arm is interesting as it implies the existence of a mechanism allowing the octopus to distinguish areas that have been searched from those that have not. This mechanism could be based on the afferent profile of mechanical novelty and other relevant cues, such as odorants, and how this profile diminishes with prolonged exposure to unchanging input. The motor response to this could take the form of lateral motion by the suckers crawling across the surface, a smaller subset of proximal suckers orienting in a new direction and causing a recruitment signal to reposition the arm, or a simple arm extension (i.e. reaching). Extension seems ineffective in the execution of a systematic search strategy as it would require the arm to lose contact with the surface, making the octopus lose all indication of its search progress given that arm configurations during the search attempt are not encoded proprioceptively.

Our model further differs from reality in that it does not take into account chemical cues. Chemical cues could inform a more directed search by the arm when near the reward, however, these were not simulated within our model. Instead, if the model comes near the reward, it is not guided by these local sensory cues. Meanwhile, the octopus’s suckers are densely innervated with chemoreceptors that may be picking up both the chemical cues from a nearby food source and its relative direction (Fouke & Rhodes, 2020; Walderon et al., 2011; Chase & Wells, 1986). Both the lack of this available information to the model and its inability to conduct systematic search patterns may account for the difference in performance between the simulated and observed search efficiency when rewards are located on the side of the task space. Another possible mechanism that combines these features could be a systematic search informed by chemical cues indicating the areas the arm has already investigated.

We attribute the difference in performance between the two observed conditions to a few primary factors. As the octopus searched the task space, the one definite location where the arm was making contact with surface features was where it was reaching through the entrance, and from this point of contact we believe a strong recruitment signal was being sent distally. Though the middle reward was secured to a surface, this surface was separated from the entrance by a concave edge oriented orthogonally to the direction of the reward relative to the entrance. The proximal-to-distal recruitment signal, likely guided by this edge, oriented the arm away from the middle and toward the side. The recurring kinematic profiles appearing in the arm speed, curvature, and wall proximity time plots reflected the prevalence of this behavior. Because of these factors, the most likely way for the arm to find the middle reward was when proximal-to-distal recruitment caused the distal arm to sweep past the reward while switching sides of the task space.

Distal-to-proximal recruitment was observed, though it usually occurred only after distal suckers found the food reward. Free proximal suckers were then “reeled in” toward the reward (see Supplemental Video 2). This was not included in the analysis primarily because this behavior was difficult to track with DeepLabCut. This behavior indicates that while distal suckers are limited in their ability to capture and manipulate prey because of their size, they may serve as scouts by locating prey then recruiting proximal suckers to capture it.

The extent to which morphology and neuroanatomical organization is conserved across units of the sucker and its adjacent length of the arm’s nerve cord and musculature has not been fully characterized. It is therefore informative to ascertain the degree to which the octopus arm’s behavior was simulated in a model whose segments were controlled by identical routines. While it has been shown that in locomotion, separate functional roles tend to be adopted by different lengths of the arms (Levy et al., 2015; Mather, 1998; Hooper, 2015; Levy & Hochner, 2017), it seems that in the context of search behavior (and possibly other behaviors employing similar recruitment patterns), sucker-arm units may be functionally identical.

By reflexively conforming its arm to the shape of the surrounding surfaces to guide its movement, these surfaces can confine the enormous degrees of freedom of the arms to a more manageable range of configurations. Serving as a lower dimensional reference during behavior, surfaces, especially sharp concave features, appear to act as coastlines in the arm’s configuration space and are perhaps used in an analogous way as coastal navigation is used by ships (Roy et al., 1999).

This behavioral strategy based on contact with the environment is interesting, as in the field of robotics collision with environmental surfaces is generally avoided. For the soft-bodied octopus arm, rather than being avoided, collision appears to be exploited as a control strategy. However, unlike the simple architecture of our task, the surface features of the octopus’s environment are convoluted and complex. The next step for this paradigm will be to investigate how this strategy is employed with tasks more closely representing this kind of surface complexity.

We believe that the number of trials used for this investigation was not sufficient to conclusively interpret the lack of improvement in task performance. However, in the event that learning is possible within this series of trials, we suggest that each condition faces its own barrier for improvement. The middle task showed no improvement because, without the arm using a surface to guide its search strategy, the brain would need to encode and recall the shape of the arms when they found the reward. However, this level of detail, namely relative position of the arm and suckers, is evidently not represented within the brain and therefore cannot be recalled to find the reward more efficiently over repeated trials. Meanwhile, the lack of improvement for the side task may represent that, as a reflex, surface conformation as initiated by sucker recruitment is limited in its capacity for improvement. However, as we mention above, there is at present a limited amount of evidence for a lack of learning; we expect that further work might provide additional evidence.

Additional limitations we would like to highlight are the lack of available females (since all animals were wild-caught, sex of animals included in the study was not under our control), and our inability to distinguish between arms with this task design (with the exception of the third right arm in males, due to the presence of the hectocotylus). The questions that we are unable to address due to this latter limitation are potential arm preference, individual arm performance, and individual arm improvement. With the results presented by Bowers et al. (2021), suggesting a possible peripheral mechanism for memory encoding within the arms of the dwarf cuttlefish *Sepia bandensis*, these are clearly interesting and viable research questions that this paradigm should seek to address moving forward. Efforts for future work will therefore be made to modify task design to be able to distinguish arms.

The paradigm we used for this investigation involved using CAD software to design and 3D print a task space that encouraged a specific behavior in the octopus. We then tracked the arm to characterize its behavioral patterns and kinematics, and developed a model of the arm equipped with mechanisms meant to generate the behaviors that we observed. Using the CAD file for the task, we were then able to simulate the experimental conditions for the model and compare the results to those of the experimental data, thus validating the mechanisms we provided the model as likely mechanisms within the octopus’s arm. This paradigm presents an effective pipeline for both characterizing underlying mechanisms of octopus arm behaviors and testing existing models of the arm. On the one hand, tasks can be designed to encourage behaviors such as search, manipulation, and locomotion, from which mechanisms can be isolated and programmed into models subjected to a simulated task in order to compare results. Alternatively, arm models can be provided with a task to predict what the octopus arm will do in a given situation, which can then be validated with a 3D printed version of the task presented to an octopus. This will be particularly useful as more sophisticated models are developed with multisensory capabilities and more accurate morphological and biomechanical properties. The fact that these tasks are printed from CAD files means that, provided printer settings are identical, the exact same task can be printed in multiple locations and used for multiple species, ensuring optimal replicability. This will necessarily be subject to slight differences in resolution depending on the capacity of the printers.

Sucker recruitment presents a simple peripheral behavioral mechanism that can lead to a number of adaptive advantages. By orienting suckers toward relevant stimuli, recruitment signals can act as an effective capture strategy. Multiple suckers can be recruited in immobilizing and handling prey where a single sucker may be insufficient. Given the minimal bandwidth between the brain and arms and the level of abstraction of mechanical information from the sucker disks (Wells & Wells, 1957), it seems like any one sucker’s ability to communicate with the brain is limited. By recruiting their neighbors toward a relevant stimulus, the representation of a stimulus within the brain can be compounded through the collective sensory fields and afferent pathways of recruited suckers. Sucker recruitment can thereby serve as an adaptive, locally controlled sensory filter. Additionally, as supported by our results, sucker recruitment serves as an effective search strategy by allowing the arm to conform to the shape of surface features, and a mechanism by which the octopus can exploit these surface features as a means to shape its arms with minimal feedback to the brain, and thereby confine the arms’ degrees of freedom to a more tractable range.

## Data availability statement

The data that support the findings of this study are available upon reasonable request from the authors. Task CAD files are also available upon request.

---

**Supplemental Figure 1.**
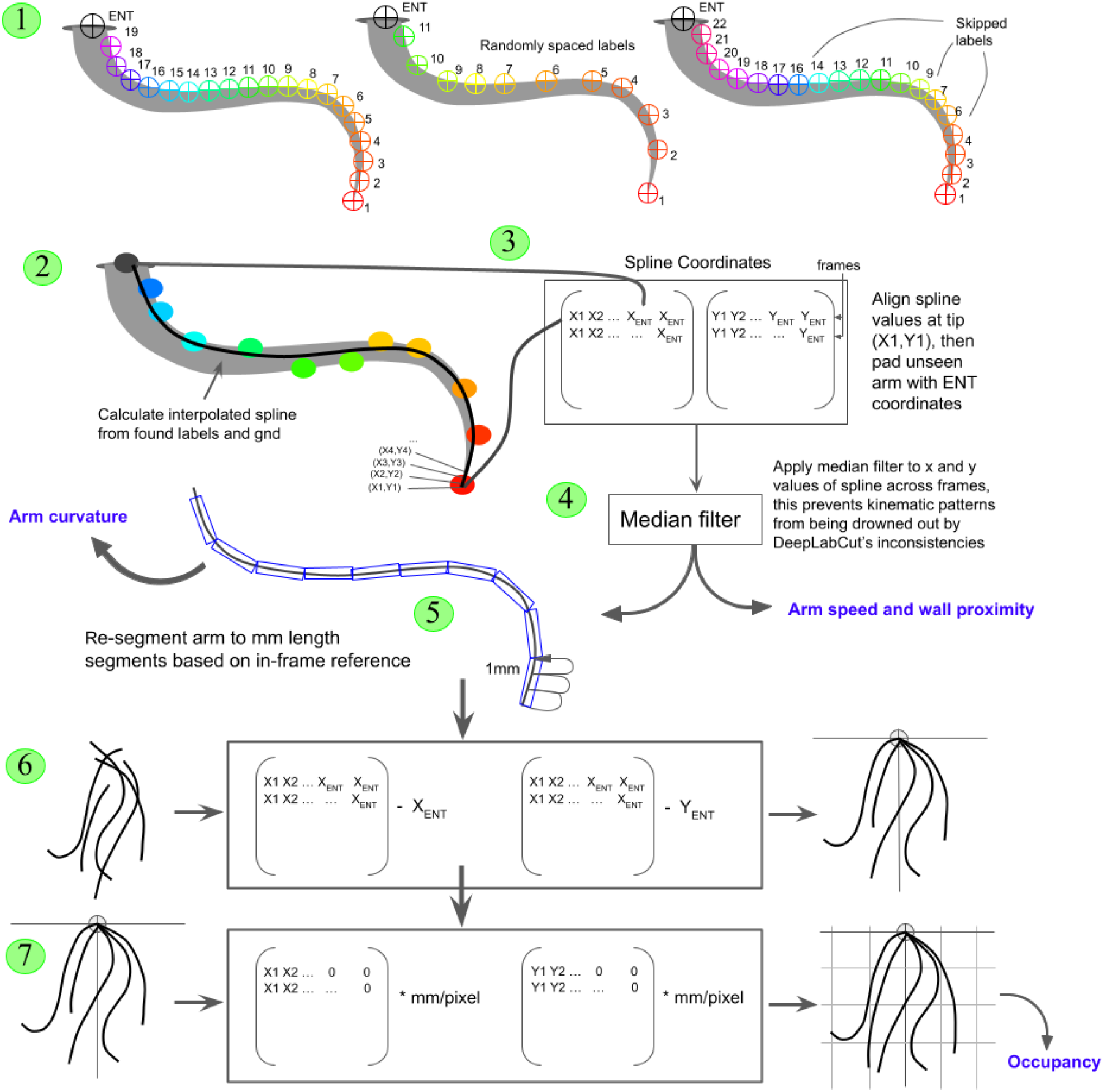
Labeling and analysis pipeline. 1) DeepLabCut pose estimation software (Mathis et al., 2018) was used to track the arm during trials. During network training, the arm was labeled with a maximum of 50 labels along its edge opposite the suckers. Labeling started at the tip with “1” and continued until the last visible section of the arm was labeled. The basic labeling pattern was consecutive and evenly spaced, however labels were occasionally spaced apart at random intervals and skipped. A separate network was used to track task features, including the entrance. 2) Labels from the DeepLabCut analysis, including the entrance label (ENT) were used to generate an interpolated spline. 3) The x and y values of each pixel of the spline were aligned within separate arrays, such that the arm tip coordinates were all represented by column 1, and each row represented frame number. The rows were padded to be equal in length using the ENT coordinates. 4) A median filter was applied to x and y coordinates of the spline across frames, preventing kinematic patterns from being drowned out by the network’s inconsistencies with the high frame rate (250 FPS) at which videos were recorded. 5) Using a tracked task feature of known length, the arm spline was re-segmented such that each segment was equal to 1mm. 6) Each arm’s ENT coordinates were subtracted from these new segment coordinates, centering the arms on the entrance locations. 7) These coordinates were scaled to a common axis using a tracked task feature of known length.

