## Supplementary figures and images for "Mechanisms of octopus arm search behavior without visual feedback"

### Supplemental Figure 1

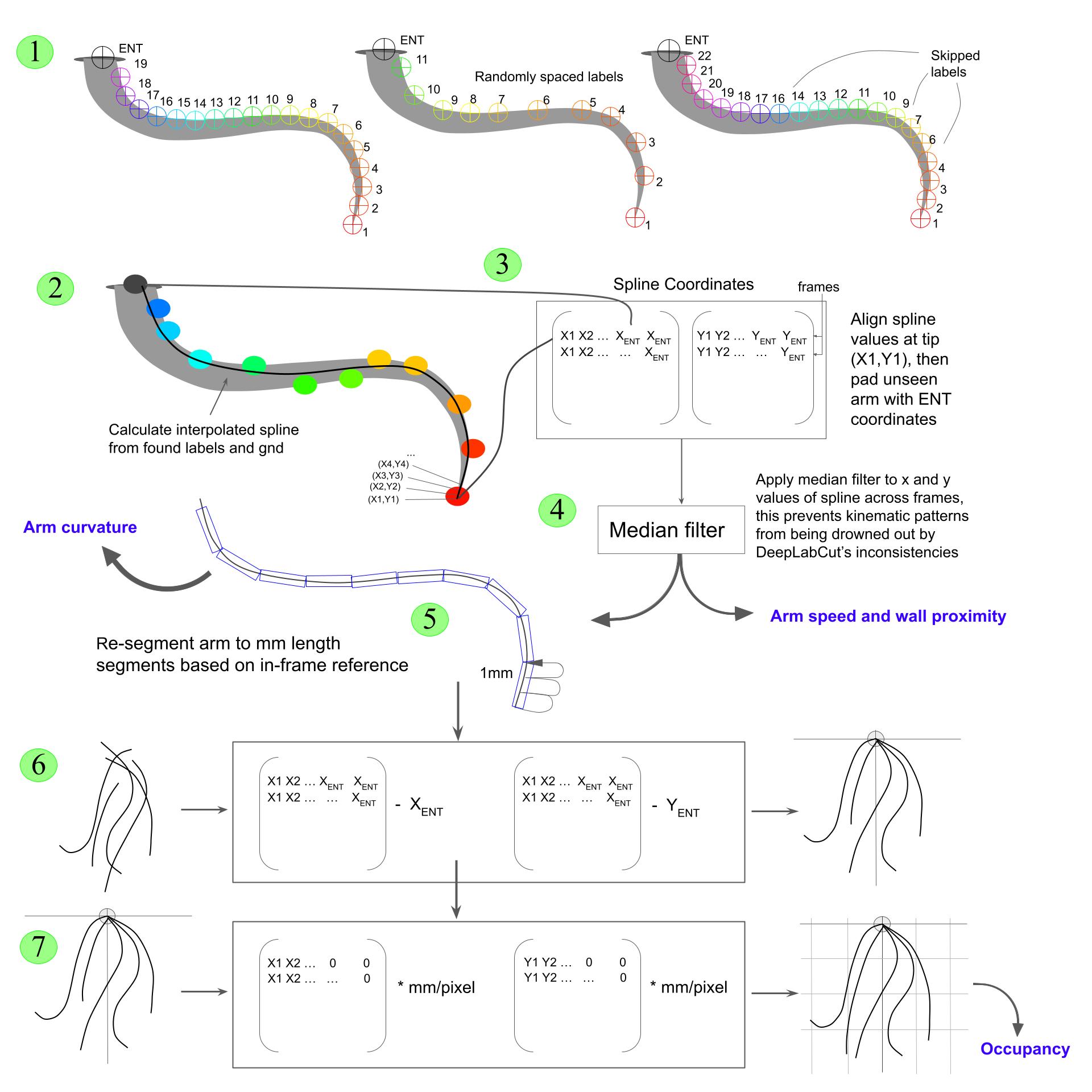
